# Multipartite mitochondrial genomes and phylogenetic relationships of five newly characterized chewing-louse species of placental mammals (Parvorder Trichodectera)

**DOI:** 10.64898/2026.10.14.759280

**Authors:** Theresa A. Spradling, Emmanuel Anane, Morgan Mescher, Grace P. Mraz, John P. Lehmkuhl, James W. Demastes

## Abstract

Fragmented mitochondrial genomes, in which genes are distributed across multiple minichromosomes rather than a single circular molecule, have evolved multiple times in parasitic lice (Psocodea). Despite this complex history, mitochondrial genomes have been characterized for only six of the roughly 400 trichodecteran louse species. Here we used PCR, cloning, and Sanger sequencing methods to increase the number of species characterized in this parvorder by determining the complete mitochondrial genomes of five additional trichodecteran lice: *Geomydoecus geomydis*, *Geomydoecus centralis*, *Thomomydoecus minor*, *Neotrichodectes mephitidis*, and *Stachiella octomaculatus*. Each genome was made up of 12–14 circular minichromosomes, consistent with the range reported in related louse lineages. Gene content and chromosome size were largely conserved among close relatives, though tRNA genes showed more mobility than did protein-coding or rRNA genes. The *nad3* gene occurred in multiple genomic positions across species. All chromosomes carried one or two conserved sequence blocks in their non-gene regions. These blocks were highly similar within each species but too divergent for alignment between species, a pattern consistent with ongoing recombination among chromosomes within each lineage. We identified additional signatures of interchromosomal recombination, including shared non-coding sequence among specific chromosome subsets in *N. mephitidis*, nearly identical *trnL1*/*trnL2* sequences in several species, and identical sequences embedded in the *cox1* and *nad2* genes. A phylogenetic analysis based on 12 mitochondrial genes resolved relationships among trichodecteran species with strong support. Carnivoran-associated lice were paraphyletic, with *N. mephitidis* being more closely related to pocket gopher lice than to other carnivoran lice, corroborating independent evidence for an ancient host switch between carnivorans and rodents. These findings expand the taxonomic sampling of fragmented mitochondrial genomes and provide further evidence that recombination actively shapes both coding and non-coding regions of these unusual genomes.

## Introduction

Mitochondrial genomes of bilaterian animals are typically compact, circular molecules of roughly 16 kb that carry the same 37 genes: 13 protein-coding genes, two ribosomal RNAs, and 22 transfer RNAs [1–3]. A minority of animal species depart from this arrangement, with the mitochondrial genome fragmented among two or more chromosomes. Some nematodes have multipartite mitochondrial genomes composed of a small number of minicircles [4] as do some rotifers [5]. Fragmented mitochondrial genomes are also observed in two of the three hemipteroid orders, Thysanoptera (thrips)[6] and Psocodea (bark lice, book lice, and parasitic lice) [7], where the condition appears to have evolved numerous times.

Our most complete understanding of the phylogeny of the Psocodea comes from de Moya et al. [8], who recognize five parvorders of parasitic lice in the infraorder Phthiraptera. Within the Psocodea, fragmentation of the mitochondrial genome was first discovered in the human body louse, *Pediculus humanus* (parvorder Anoplura), which has a mitochondrial genome that is subdivided into 18 minichromosomes, each with only one to three genes per chromosome [9]. Numerous parasitic and non-parasitic lice have since been shown to have fragmented mitochondrial genomes. Fragmentation has now been observed in all five parvorders of parasitic lice, including the Trichodectera [10], Anoplura [11–20], Rhynchophthirina [18], Ischnocera [21,22], and Amblycera [23,24].

Whereas the onset of fragmentation has been determined to have occurred multiple times within the parvorders Ischnocera and Amblycera [7,21–24], based on current evidence, fragmentation presumably arose once in the common ancestor of Anoplura, Rhynchophthirina, and Trichodectera, which form a well-supported clade [8], and has been maintained in these lineages since. Fragmentation also occurred more than once in *Liposcelis* [7], a non-parasitic genus of book lice that de Moya et al. [8] place in the infraorder Liposcelidetae.

Fragmentation is correlated with several distinctive genomic features. For example, fragmented mitochondrial genomes have shown faster sequence evolution and more gene rearrangement [7], and in a matched-pair analysis of parasitic lice, fragmented lineages had significantly longer mitochondrial branch lengths and more rearranged gene orders than their single-chromosome relatives [24]. It remains unclear whether an elevated mutation rate drives genomic fragmentation, whether fragmentation increases the mutation rate, or whether both arise from a common underlying cause [7,24]. Fragmented genomes also tend to show a weaker AT bias than closely related species with a single circular chromosome [8]. Heteroplasmy has been reported in fragmented genomes [25], but it appears to be common in both fragmented and non-fragmented mitochondrial genomes [26]. Fragmentation may also affect gene expression. Dunn and Shao [27] recently documented highly variable transcription rates among mitochondrial rRNA and protein-coding genes in fragmented genomes. This is plausible because each chromosome can be transcribed independently, unlike single-chromosome genomes, in which transcription is initiated regularly at only three sites. Because fragmentation has arisen many times in lice, Dunn and Shao [27] suggest that transcription may proceed differently in different lineages.

The cause of fragmentation remains unresolved. One explanation is that relaxed selective constraint, combined with small effective population size, allows fragmentation to become established, with recombination among chromosomes as a byproduct of matching sequences among chromosomes [9,22,24,28]. Attempts at determining a molecular mechanism behind fragmentation have revealed further complexity. Because some lice lack a functional mitochondrial single-stranded DNA-binding protein gene (*mtSSB*) in the nuclear genome, its loss was proposed as a potential driver of fragmentation [28]. However, N. Song and Shao [29] found through increased sampling that, although fragmented species often lack *mtSSB*, some species with fragmented mitochondrial genomes retain an apparently functional copy. This suggests that loss of *mtSSB* may be a consequence of its reduced utility in fragmented genomes rather than a direct cause of fragmentation [28,30].

Given the multiple origins of fragmentation in mitochondrial genomes and the possibility of lineage-specific transcriptional dynamics [27], a better accounting of how many fragmented genomes exist, and of the characteristics of each, is needed. Sweet et al. [22] noted that further study of louse taxa should help clarify the process and direction of fragmentation. Here we examine lice from the parvorder Trichodectera, which include approximately 400 described species [31]. Of these, the mitochondrial genomes of only six species have been characterized. *Geomydoecus aurei* occurs on the pocket gopher *Thomomys bottae* (Rodentia: Geomyidae), and its mitochondrial genome has been published [32,33]. Four additional species of trichodecteran mitochondrial genomes were characterized by Song et al. [10]: *Trichodectes canis*, a parasite of dogs, and *Bovicola bovis*, *B. ovis*, and *B. caprae*, which parasitize cattle, sheep, and goats, respectively. Ponce et al. recently characterized the mitochondrial genome of *Trichodectes melis*, the parasite of the European badger, *Meles meles* [34].

In this study, we determined the mitochondrial genomes of five species, including two congeners of *G. aurei*. *Geomydoecus centralis* occurs on *T. bottae* in the same region of New Mexico as *G. aurei*, and the two species sometimes co-occur near a host hybrid zone [35,36]. *Geomydoecus geomydis* parasitizes the pocket gopher *Geomys bursarius* in Iowa [31,37]. *Thomomydoecus* is the sister genus to *Geomydoecus* [38], and we represented this genus by characterizing the mitochondrial genome of *Thomomydoecus minor*, which parasitizes *T. bottae* and co-occurs with both *G. centralis* and *G. aurei* [35]. Finally, we characterized the mitochondrial genomes of two louse species that occur on carnivorans: *Neotrichodectes mephitidis*, found on striped skunks (*Mephitis mephitis* [31]), and *Stachiella octomaculatus*, found on raccoons (*Procyon lotor* [31]). Our objectives were to characterize the number and content of chromosomes in each species, to compare gene and non-gene regions of chromosomes among species, examine evidence for recombination among chromosomes within a species, and examine phylogenetic relationships within the Trichodectera.

## Methods

### Specimen collection

Lice were collected from hosts either salvaged from the road or trapped under the appropriate governmental permits, ethical guidelines set by the American Society of Mammalogists [39], and with the prior approval of the University of Northern Iowa Institutional Animal Care and Use Committee.

Specimens of *Geomydoecus geomydis* were collected from the fur of three plains pocket gophers (*Geomys bursarius*; vouchers TAS 735, 746, and 747), captured in Iowa: Black Hawk County, near Cedar Hills Sand Prairie (42° 36’38” N, 92° 34’35” W), 855ft. Iowa pocket gophers were trapped with permission of the Iowa Department of Natural Resources (Code of Iowa 481A.65 and Iowa Administrative Code 571.111, permit number, SC553).

Pocket gophers (*Thomomys bottae*) were collected from Socorro County, New Mexico, with the approval of the New Mexico Department of Game and Fish (Permit #3500). Specimens of *G. centralis* were collected from the same host individual (TAS 773; New Mexico; Socorro County: Polvadera 34.18393, -106.90427). Specimens of *T. minor* were collected from a nearby host individual (TAS 761; New Mexico; Socorro County: 0.9 mi S, 0.1mi W La Joya).

Specimens of *N. mephitidis* were collected from a skunk (*Mephitis mephitis*) that was killed by a farmer (Iowa: Black Hawk County, 6 mi. W Dunkerton). Specimens of *S. octomaculatus* were collected from a raccoon (*Procyon lotor*) that had been killed on the road (Iowa: Butler County, 3 mi NW New Hartford on C57). Both hosts were collected under Iowa Department of Natural Resources Permit SC553.

### DNA extraction

Individual lice were placed on microscope slides, which were placed on top of a frozen block. With the aid of a dissecting microscope, the louse’s abdomen was punctured four times using a #2 insect pin [33,40]. Genomic DNA was extracted from individual lice using Thermo Scientific Gene JET Genomic DNA purification Kit following manufacturer’s recommendations with two exceptions, one being a longer digestion with proteinase K solution at 56°C for 28 hours to allow more lysis of insect cells. Also, DNA was eluted off the kit’s columns in the final step using only 40 µl of water due to the small sizes of these insects. Cleared louse bodies, consisting of exoskeletons and durable internal structures, were removed from the extraction following proteinase K digestion and mounted on microscope slides to serve as voucher specimens.

### Chromosomal architecture

Nucleotide sequence and chromosomal content of all five mitochondrial genomes were determined using the same approach with a set of common techniques that included Polymerase Chain Reaction (PCR), Sanger Sequencing, and TA cloning. All polymerase chain reactions were performed using 1.0 µl template DNA in a 10-20 µl reaction with HotStart GoTaq Green Master Mix (Promega, Madison, WI) and thermal-cycling parameters appropriate to the primer pair. Negative controls were included to ensure reaction components were free from DNA contaminants. The amplified products were verified on 0.8% agarose gels. All PCR products generated for nucleotide sequence analysis were treated with ExoSap-IT (USB, Cleveland, OH) and sequenced directly using Sanger sequencing at the Iowa State University DNA Facility. Sequenced products were trimmed, edited manually, and aligned to generate consensus sequences using Geneious 11.1.5 [41] such that there was at least double coverage of all regions of the chromosomes.

All chromosomes analyzed herein contained regions of heteroplasmy, so a cloning approach was used to cleanly sequence through these regions. As part of the fundamental approach of this work, cloning also was necessary to sort sequences that were generated using primers known to amplify multiple chromosomes or nuclear copies of mitochondrial genes. PCR products were cloned using the TOPO TA cloning Kit with the PCR 4-TOPO vector with Invitrogen TOP-10 *Escherichia coli* competent cells (Life Technologies Carlsbad, CA). Transformed cells were spread on LB-agar selective plates with either kanamycin or ampicillin and allowed to grow overnight at 37°C. Colonies were picked and amplified using primers that targeted plasmid regions surrounding the PCR insert. The amplified products were sent for Sanger sequencing, and in each case, multiple clones were sequenced to assess heteroplasmy.

Genomic analysis of each species began with PCR using universal primers for short regions of the *cox1* and *rrnL* genes. The universal *cox1* primers LCO1490 and HCO2198 [42] amplified an approximately 700 base-pair (bp) region of the mitochondrial genome. For *G. centralis*, universal primers H7005 and L6625 [43] were used to amplify a 379 bp region of *cox1* instead of the larger fragment of the gene. Similarly, a 350 bp region of the *rrnL* gene was targeted using universal *rrnL* PCR primers (Lx16SF and Lx16SR) [12,19]. These products were either sequenced directly or cloned and then sequenced to yield short, unambiguous mitochondrial sequences for the *cox1* and *rrnL* gene. Cloning was necessary when nuclear copies of these genes coamplified with mitochondrial copies, as they did in both *G. geomydis* (*cox1*) and *T. minor* (*rrnL*).

The short mitochondrial *cox1* and *rrnL* sequences were used to design primers that faced outward on their respective chromosomes (S1-S5 Tables). Repeated failure of well-designed primers to amplify a circular chromosome was taken as evidence that a linear (i.e., nuclear) copy of the gene had been amplified, and those sequences were eliminated from further analysis. Amplification products from apparently circular chromosomes were cloned and sequenced as described above to yield complete *cox1* and *rrnL* chromosomes for each species with clear sequences through regions of both length and nucleotide-sequence heteroplasmy. These products confirmed the circular structure of the chromosome.

Alignment of the noncoding regions of the *cox1* and *rrnL* chromosomes was performed to detect nucleotide sequences that were conserved between the two chromosomes for each of the five species analyzed. These common sequences were then used to design “multi-circle” primers (S1-S5 Tables) that targeted the common sequence region of other mitochondrial minichromosomes of the genome. Multi-circle primers faced “outward” on the circle to amplify a nearly complete chromosome ranging from approximately 1,200 bp to 2,100 bp depending on the chromosome and the species. Cloning was then used to individually analyze sequences for each chromosome amplified by the multi-circle PCR products of each species. Hundreds of multi-circle clones were analyzed for each species until the full complement of minichromosomes was determined multiple times.

Circularization of the nucleotide sequence of each chromosome was completed by designing an additional set of primers that faced away from one another at positions within the protein-coding gene of each species (S1-S5 Tables). Amplifications using these primers, followed by cloning and sequencing reactions, yielded additional coverage of each chromosome, including the region spanning the multi-circle primer sites and the region internal to them.

### Mitochondrial genome annotation and accession

Annotation was performed using MITOS2 software [44] on the European Galaxy server, a public Galaxy instance hosted by the European Galaxy project [45,46]. MITOS2 (Mitochondrial Genome Annotation Server 2) relies on Hidden Markov Models (HMMS) tailored for mitochondrial gene profiles to annotate genomic regions accurately [44,47]. MITOS2 is a widely used tool for annotating and analyzing mitochondrial genomes from various organisms using the result of BLASTx searches against the amino acid sequences of the annotated proteins of metazoan mitochondrial genomes [44]. Similarly, transfer RNA genes were predicted by MITOS2 using their characteristic secondary structures [48]. The secondary structure of tRNA genes was further verified on tRNAscan-SE 2.0 [49,50] and/or ARAGORN (Version 1.2.41) [48] run on the European Galaxy server [45,46].

Complete, annotated chromosome sequences were archived in Genbank: *G. geomydis* PZ162147-PZ162158, *G. centralis* PP798382-PP798391, *T. minor* PQ469563-PQ469574, *N. mephitidis* PZ094309-PZ094322, *S. octomaculatus* PZ137458-PZ137470. Chromosome maps (Figs 1-5) were visualized for linear presentation using SnapGene® software (www.snapgene.com; version 8.2.2).

**Fig 1.**
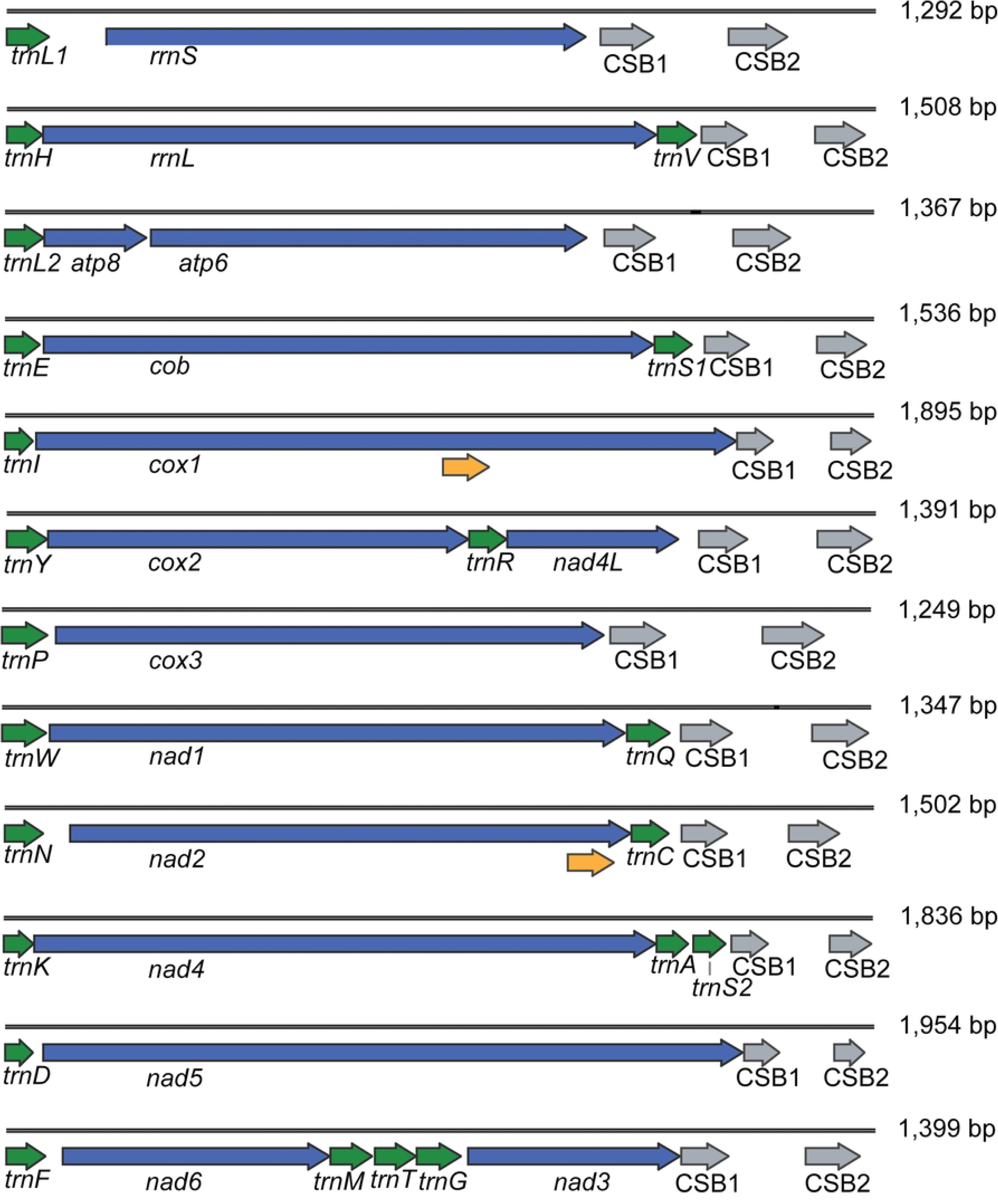
Twelve circular mitochondrial chromosomes of *Geomydoecus geomydis*, linearized for display purposes. Gene name and transcription orientation are given along with the location of Conserved Sequence Blocks (CSBs). Yellow arrows indicate the locations of nucleotide sequence that are identical on *cox1* and *nad2*. Numbers at the right indicate the length of each chromosome (bp).

### Non-gene region alignment and determination of CSBs

To identify sequences common to all chromosomes (CSBs), the non-gene regions of all mitochondrial chromosomes were extracted and annotated using Geneious 11.1.5 [41]. Multiple sequence alignment was performed in Geneious using a “map to reference” approach. Alignments were visually inspected and subsequently defined using strong percent-pairwise identity across the alignments coupled with large numbers of nucleotide sites that were identical among chromosomes.

### Phylogenetic analysis

Nucleotide sequences from the protein-coding and rRNA genes of ten trichodecteran species (representing trichodecteran lice available in Genbank on May 6, 2026) and the outgroup, *Pediculus humanus corporis* [9], were aligned in Geneious 11.1.5 [41] using the MAFFT v7.450 plugin [51,52]. Alignments for the two rRNA genes were performed with the L-INS-i algorithm for local pairwise alignment, appropriate for sequences with conserved regions surrounded by long gaps. Alignments for the protein-coding genes were performed using the G-INS-i algorithm, appropriate for globally alignable sequences. These alignments were passed through the TrimAl program (version 1.5.1), and treated with the GappyOut protocol, all run on the European Galaxy server [45,46,53]. The trimmed files were again visually inspected and trimmed at the beginning and end of genes where necessary to produce gene alignments of equal lengths for all species. These 12 trimmed gene alignments were then concatenated in Geneious 11.1.5 for phylogenetic analysis.

The concatenated data set was uploaded to the European Galaxy server [45,46], where partitioned maximum likelihood analysis was performed with IQ-TREE (Version 2.4.0) using the edge-unlinked proportional partition model [54,55]. In this analysis, ModelFinder was used to calculate the best substitution model for each gene partition [55,56], and each gene received its own independent set of branch lengths. IQ-TREE was also used to perform 10,000 non-parametric bootstrap replicates.

## Results

### Chromosomal structure, size, and gene content

Mitochondrial genomes of the five newly described trichodecteran species each comprised 12–14 circular minichromosomes (Table 1, Figs 1-5). Of these species, lice that parasitize pocket gophers, *G. geomydis*, *G. centralis*, and *T. minor*, exhibited 12 minichromosomes (Figs 1-3). *Neotrichodectes mephitidis* had 14 minichromosomes (Fig 4), and *S. octomaculatus* had 13 (Fig 5).

**Fig 2.**
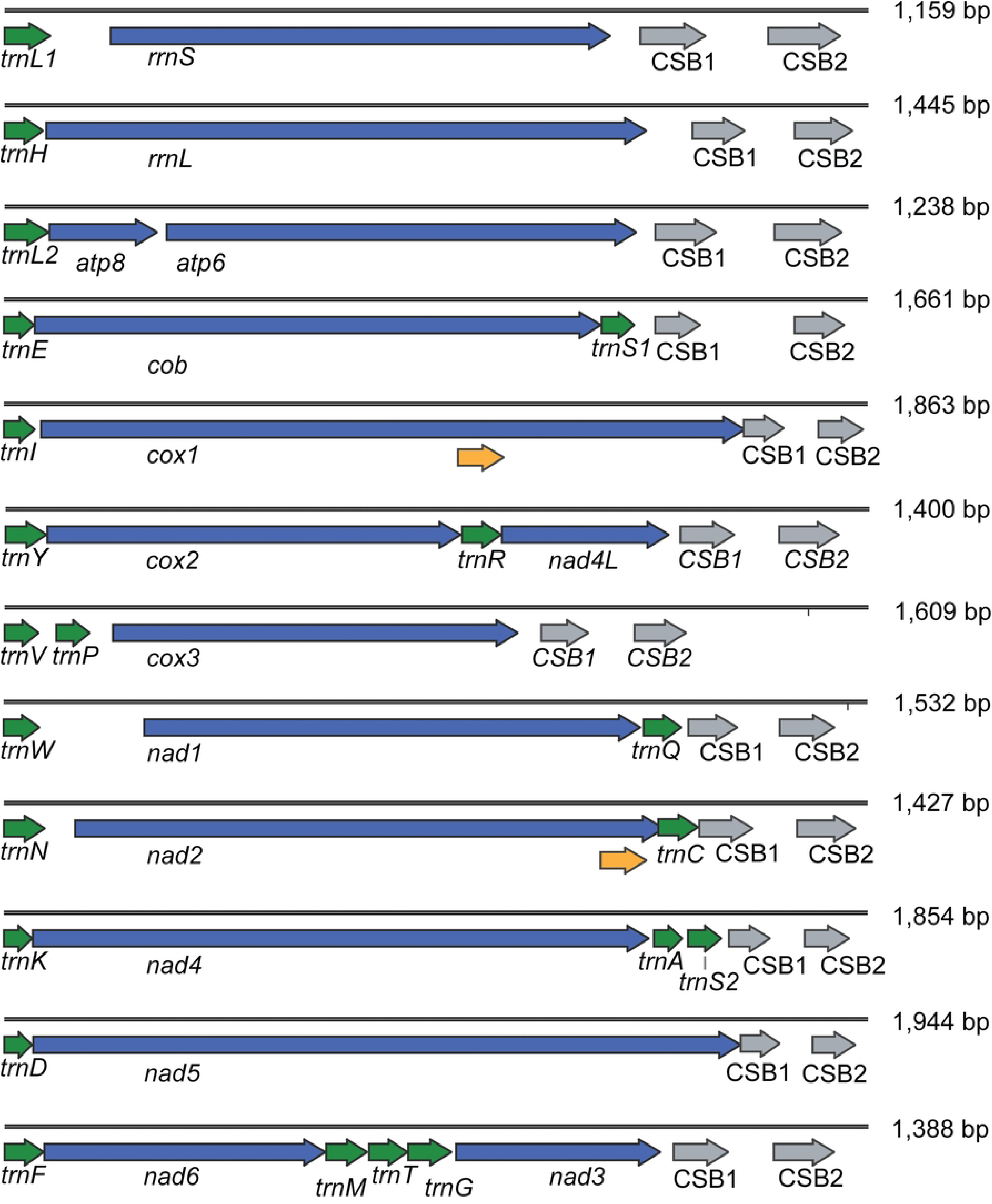
Twelve circular mitochondrial chromosomes of *Geomydoecus centralis*, linearized for display purposes. Gene name and transcription orientation are given along with the location of Conserved Sequence Blocks (CSBs). Yellow arrows indicate the locations of nucleotide sequence that are identical on *cox1* and *nad2*. Numbers at the right indicate the length of each chromosome (bp).

**Fig 3.**
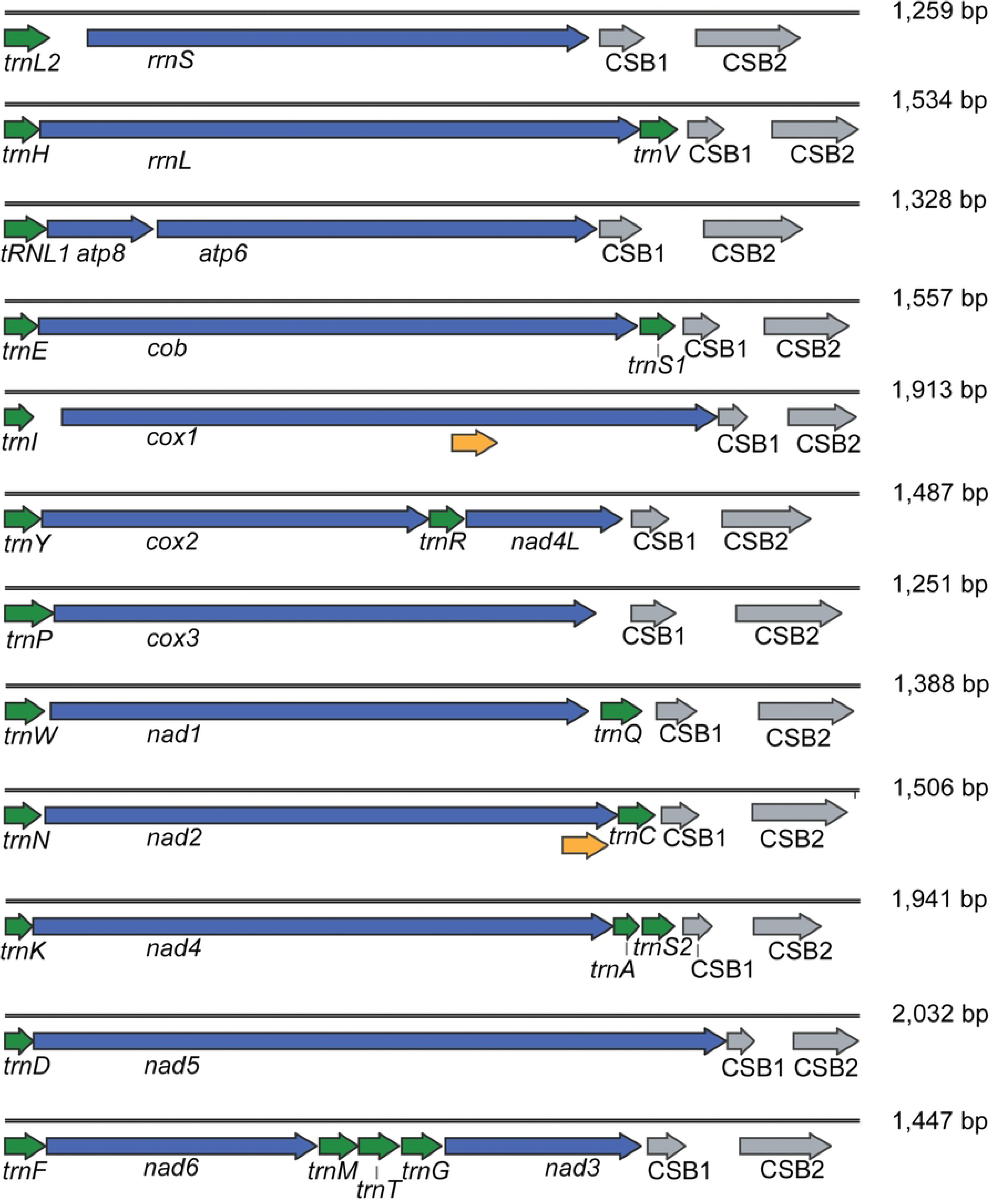
Twelve circular mitochondrial chromosomes of *Thomomydoecus minor*, linearized for display purposes. Gene name and transcription orientation are given along with the location of Conserved Sequence Blocks (CSBs). Yellow arrows indicate the locations of nucleotide sequence that are identical on *cox1* and *nad2*. Numbers at the right indicate the length of each chromosome (bp).

**Fig 4.**
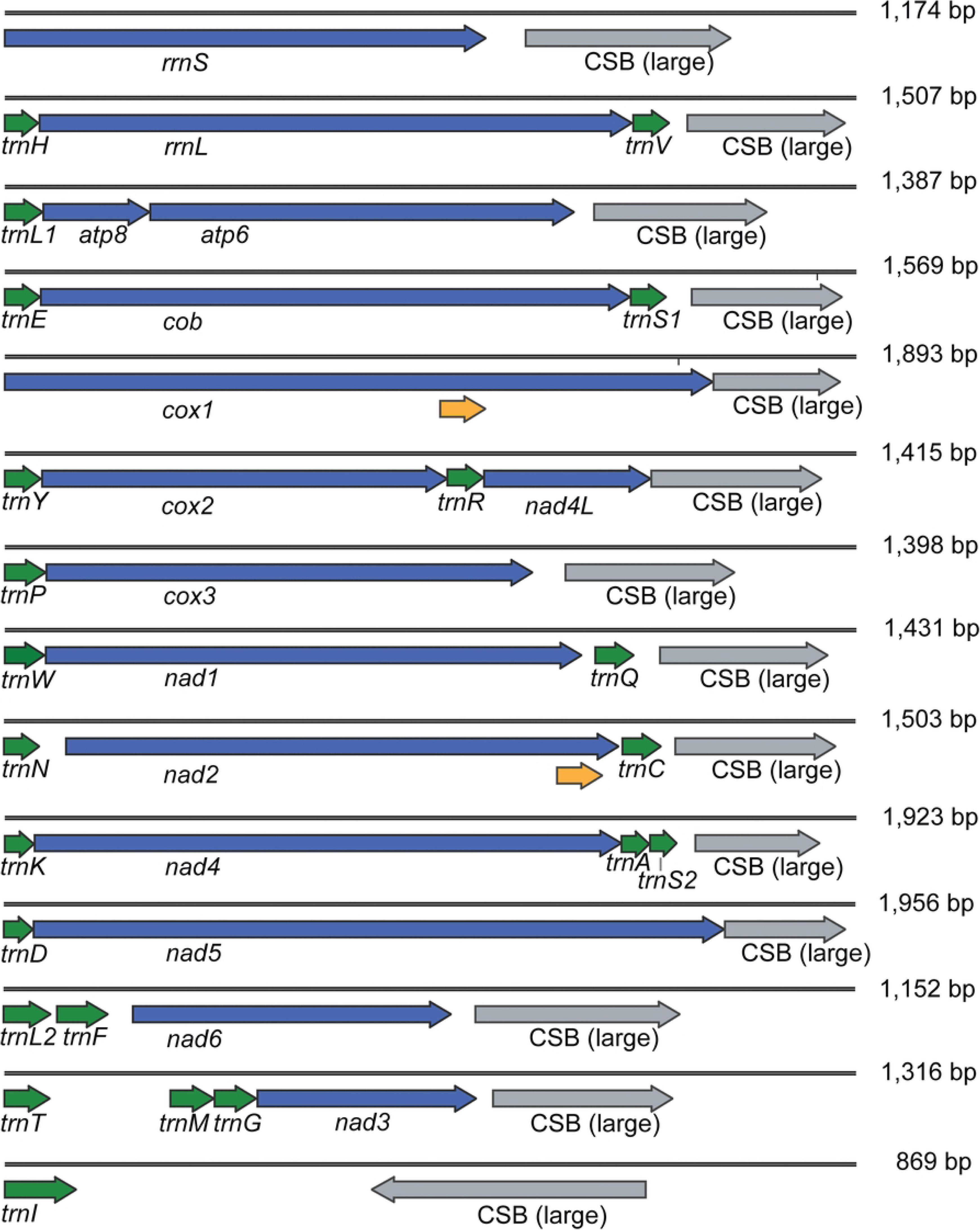
Fourteen circular mitochondrial chromosomes of *Neotrichodectes mephitidis*, linearized for display purposes. Gene name and transcription orientation are given along with the location of Conserved Sequence Blocks (CSBs). Yellow arrows indicate the locations of nucleotide sequence that are identical on *cox1* and *nad2*. Numbers at the right indicate the length of each chromosome (bp).

**Table 1:** Gene order, chromosome size, gene size, and non-gene content by chromosome for five new trichodecteran lice.

| Chromosome and Species | Chromosome Gene Order <sup>a</sup> | Size of Chromosome | Size of Protein or rRNA Genes | Non-gene Region Size <sup>b</sup> / % of Chromosome |
| --- | --- | --- | --- | --- |
| **1. *rrnS*** |  |  |  |  |
| *G. geomydis* | *L1-rrnS* | 1,292 | 715 | 513 in 2 segments / 40% |
| *G. centralis* | *L1-rrnS* | 1,159 | 673 | 423 in 2 segments / 37% |
| *T. minor* | *L2-rrnS* | 1,259 | 740 | 452 in 2 segments / 36% |
| *N. mephitis* | *rrnS* | 1,174 | 665 | 509 in 1 segment / 43% |
| *S. octomaculatus* | *rrnS* | 1,218 | 711 | 507 in 1 segment / 42% |
| **2. *rrnL*** |  |  |  |  |
| *G. geomydis* | *H-rrnL-V* | 1,508 | 1,067 | 309 in 1 segment / 20% |
| *G. centralis* | *H-rrnL* | 1,445 | 1,006 | 374 in 1 segment / 26% |
| *T. minor* | *H-rrnL-V* | 1,534 | 1,079 | 324 in 1 segment / 21% |
| *N. mephitis* | *H-rrnL-V* | 1,507 | 1,051 | 328 in 3 segments / 26% |
| *S. octomaculatus* | *H-rrnL* | 1,415 | 1,116 | 236 in 1 segment / 17% |
| **3. *atp8, atp6*** |  |  |  |  |
| *G. geomydis* | *L2-atp8-atp6* | 1,367 | 162, 687 | 348 in 2 segments / 25% |
| *G. centralis* | *L2-atp8-atp6* | 1,238 | 156, 675 | 337 in 2 segments / 27% |
| *T. minor* | *L1-atp8-atp6* | 1,328 | 165, 684 | 412 in 2 segments / 31% |
| *N. mephitis* | *L1-atp8-atp6* | 1,387 | 174, 693 | 502 in 2 segments / 36% |
| *S. octomaculatus* | *L2-atp8-atp6* | 1,259 | 168, 693 | 330 in 2 segments / 26% |
| **4. *cob*** |  |  |  |  |
| *G. geomydis* | *E-cob-S1* | 1,536 | 1,080 | 325 in 2 segments / 21% |
| *G. centralis* | *E-cob-S1* | 1,661 | 1,092 | 446 in 1 segment / 27% |
| *T. minor* | *E-cob-S1* | 1,557 | 1,092 | 339 in 3 segments / 22% |
| *N. mephitis* | *E-cob-S1* | 1,569 | 1,089 | 348 in 1 segment / 22% |
| *S. octomaculatus* | *E-cob-S1* | 1,504 | 1,101 | 267 in 1 segment / 18% |
| **5. *cox1*** |  |  |  |  |
| *G. geomydis* | *I-cox1* | 1,895 | 1,530 | 302 in 2 segments / 16% |
| *G. centralis* | *I-cox1* | 1,863 | 1,521 | 259 in 2 segments / 14% |
| *T. minor* | *I-cox1* | 1,913 | 1,470 | 306 in 2 segments / 16% |
| *N. mephitis* | *cox1* | 1,893 | 1,578 | 315 in 1 segment / 17% |
| *S. octomaculatus* | *I-cox1* | 1,839 | 1,530 | 242 in 1 segment / 13% |
| **6. *cox2, nad4L*** |  |  |  |  |
| *G. geomydis* | *Y-cox2-R-nad4L* | 1,391 | 675, 276 | 313 in 1 segment / 23% |
| *G. centralis* | *Y-cox2-R-nad4L* | 1,400 | 675, 273 | 323 in 2 segments / 23% |
| *T. minor* | *Y-cox2-R-nad4L* | 1,487 | 675, 273 | 414 in 3 segments / 28% |
| *N. mephitis* | *Y-cox2-R-nad4L* | 1,415 | 675, 279 | 340 in 2 segments / 24% |
| *S. octomaculatus* | *cox2-R-nad4L* | 1,360 | 666, 246 | 316 in 2 segments / 23% |
| **7. *cox3*** |  |  |  |  |
| *G. geomydis* | *P-cox3* | 1,249 | 789 | 393 in 2 segments / 31% |
| *G. centralis* | *V-P-cox3* | 1,609 | 756 | 724 in 3 segments / 45% |
| *T. minor* | *P-cox3* | 1,251 | 795 | 384 in 1 segment / 31% |
| *N. mephitidis* | *P-cox3* | 1,398 | 801 | 529 in 1 segment / 38% |
| *S. octomaculatus* | *L1-cox3* | 1,134 | 807 | 259 in 1 segment / 23% |
| **8. nad1** |  |  |  |  |
| *G. geomydis* | *W-nad1-Q* | 1,347 | 894 | 313 in 2 segments / 23% |
| *G. centralis* | *W-nad1-Q* | 1,532 | 882 | 556 in 3 segments / 36% |
| *T. minor* | *W-nad1-Q* | 1,388 | 876 | 381 in 3 segments / 27% |
| *N. mephitidis* | *W-nad1-Q* | 1,431 | 903 | 393 in 2 segments / 27% |
| *S. octomaculatus* | *W-nad1-Q* | 1,342 | 906 | 305 in 2 segments / 23% |
| **9. nad2** |  |  |  |  |
| *G. geomydis* | *N-nad2-C* | 1,502 | 972 | 396 in 2 segments / 26% |
| *G. centralis* | *N-nad2-C* | 1,427 | 975 | 326 in 2 segments / 23% |
| *T. minor* | *N-nad2-C* | 1,505 | 1,011 | 364 in 3 segments / 49% |
| *N. mephitidis* | *N-nad2-C* | 1,503 | 978 | 392 in 3 segments / 26% |
| *S. octomaculatus* | *N-nad2-C* | 1,492 | 990 | 369 in 3 segments / 25% |
| **10. nad4** |  |  |  |  |
| *G. geomydis* | *K-nad4-A-S2* | 1,836 | 1,317 | 316 in 2 segments / 17% |
| *G. centralis* | *K-nad4-A-S2* | 1,854 | 1,326 | 331 in 3 segments / 17% |
| *T. minor* | *K-nad4-A-S2* | 1,941 | 1,320 | 425 in 2 segments / 14% |
| *N. mephitidis* | *K-nad4-A-S2* | 1,923 | 1,329 | 402 in 1 segment / 21% |
| *S. octomaculatus* | *K-nad4-A-S2* | 1,764 | 1,320 | 240 in 2 segments / 14% |
| **11. nad5** |  |  |  |  |
| *G. geomydis* | *D-nad5* | 1,954 | 1,578 | 312 in 2 segments / 16% |
| *G. centralis* | *D-nad5* | 1,944 | 1,596 | 284 in 2 segments / 15% |
| *T. minor* | *D-nad5* | 2,032 | 1,650 | 313 in 1 segment / 15% |
| *N. mephitidis* | *D-nad5* | 1,956 | 1,590 | 297 in 1 segment / 15% |
| *S. octomaculatus* | *D-nad5* | 1,936 | 1,605 | 265 in 1 segment / 14% |
| **12. nad6** |  |  |  |  |
| *G. geomydis* | *F-nad6-M-T-G-nad3* | 1,396 | 432, (342) | 353 in 4 segments / 25% |
| *G. centralis* | *F-nad6-M-T-G-nad3* | 1,388 | 453, (333) | 307 in 4 segments / 22% |
| *T. minor* | *F-nad6-M-T-G-nad3* | 1,447 | 459, (333) | 308 in 5 segments / 21% |
| *N. mephitidis* | *L2-F-nad6<sup>c</sup>* | 1,152 | 432 | 585 in 3 segments / 51% |
| *S. octomaculatus* | *F-nad6-[T]<sup>d</sup>____-M-G-nad<sup>e</sup>* | 1,579 | 496, (381) | 432 in 1 segment / 27% |
| **13. nad3-only** |  |  |  |  |
| *G. geomydis* | None observed | None | None | None |
| *G. centralis* | None observed | None | None | None |
| *T. minor* | None observed | None | None | None |
| *N. mephitidis* | *T-____-M-G-nad3<sup>e</sup>* | 1,316 | 339 | 772 in 2 segments / 58% |
| *S. octomaculatus* | None observed | None | None | None |
| **tRNA-only** |  |  |  |  |
| *G. geomydis* | None observed | None | None | None |
| *G. centralis* | None observed | None | None | None |
| *T. minor* | None observed | None | None | None |
| *N. mephitis* | *I* | 869 | None | 795 in 1 segment / 91% |
| *S. octomaculatus* | *Y-V-P* | 873 | None | 664 in 3 segments / 76% |

<sup>a</sup> tRNA genes are listed by their single letter code.
<sup>b</sup> "non-gene region" excludes protein-coding, rRNA, and tRNA genes, which are presumed functional
<sup>c</sup> *nad3* is located on a 13<sup>th</sup> chromosome in this species
<sup>d</sup> [T] represents the *trnT* gene transcribed in the opposite direction than other genes
<sup>e</sup> \_\_\_\_ represents a long region of non-gene sequence between other genes

When sorted by the major protein-coding or rRNA genes that each chromosome carried, the total size of chromosomes was often similar among species, although there were some exceptions. For example, the *cox3* chromosome was larger in *G. centralis* than in the other species studied, which was the result of differences in the non-gene content of the chromosome and the addition of a tRNA gene (Table 1).

Most chromosomes carried one major gene (either an rRNA gene or a protein-coding gene) and one to four tRNA genes. Two chromosomes carried a combination of two relatively small protein-coding genes (*atp8* with *atp6* and *cox2* with *nad4L*). The *nad3* gene was the only gene found in alternative chromosomal locations in the newly studied species. In *N. mephitidis*, *nad3* was located alone on a chromosome (Fig 4). In the other four species, *nad3* was found on the *nad6* chromosome; however, the relative position of *nad3* to *nad6* differed between species. In the lice of pocket gophers, *nad3* was located near *nad6* (Figs 1-3), but in *S. octomaculatus* the large non-coding region was found between the two genes (Fig 5). Both *N. mephitidis* and *S*. *octomaculatus* had one chromosome carrying only tRNA genes: *trnI* in *N. mephitidis*, and *trnY*, *trnV*, and *trnP* in *S. octomaculatus* (Table 1).

**Fig 5.**
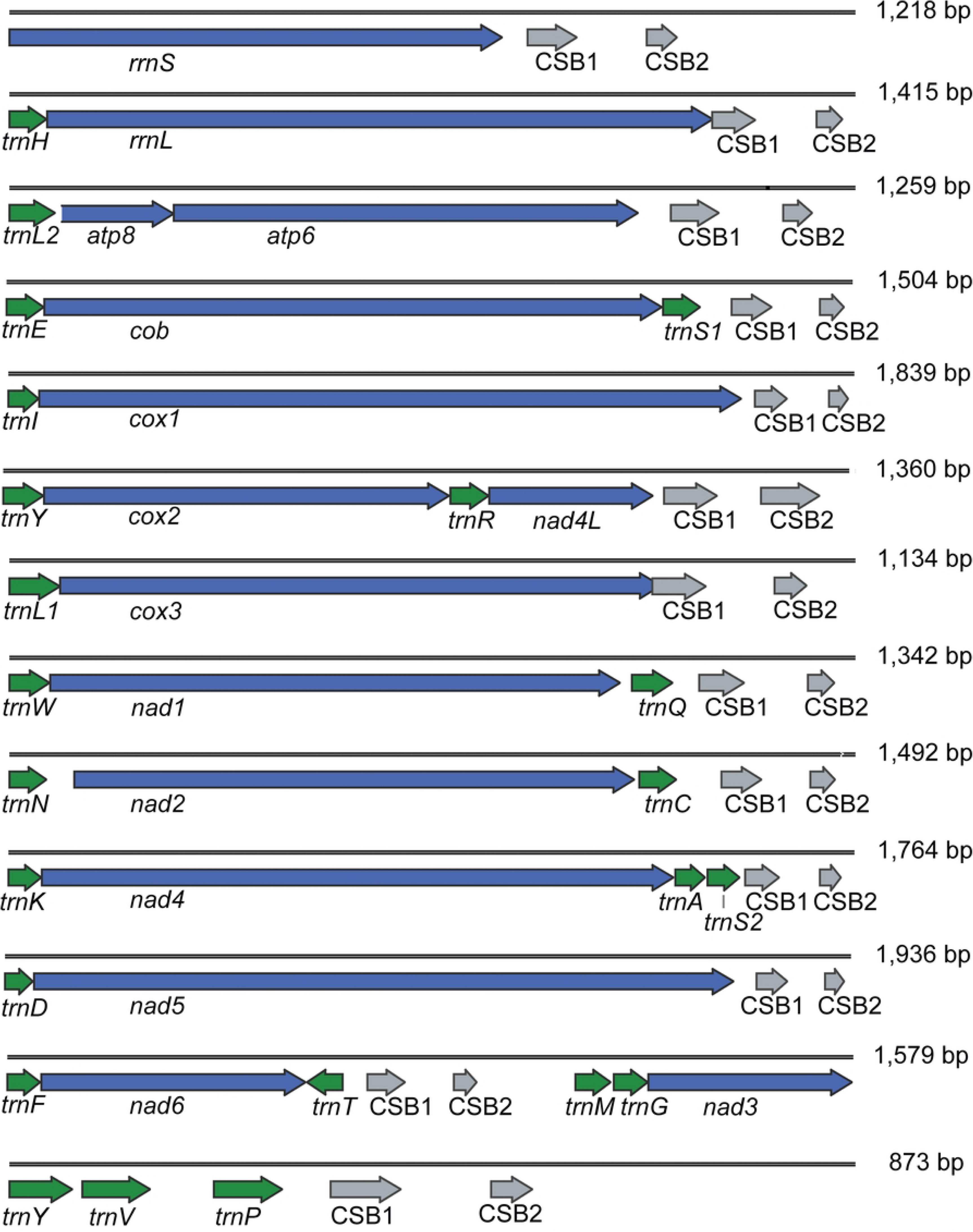
Thirteen circular mitochondrial chromosomes of *Stachiella octomaculatus*, linearized for display purposes. Gene name and transcription orientation are given along with the location of Conserved Sequence Blocks (CSBs). Numbers at the right indicate the length of each chromosome (bp).

No pseudogenes were discovered in these species, and there were no instances of any gene being found on multiple chromosomes. Moreover, all genes were oriented for transcription in the same direction as each other except for *trnT* in *S. octomaculatus*, which had a reverse orientation (Table 1, Figs 1-5).

### Shared elements among chromosomes

Each species had a non-gene region of several hundred bp, containing no tRNA, rRNA, or protein-coding sequence ("non-gene" regions; Table 1). Within the non-gene region, the newly characterized genomes exhibited one large subregion, or two small subregions, representing CSBs that exhibited 94% or greater percent-pairwise identity across the alignments coupled with over 80% nucleotide sites that were identical among chromosomes (Table 2). CSBs of the 5 species we studied were differentiated from each other enough that sequence alignments could not be created with confidence. Each of the genomes we studied contained within them GC-rich regions and AT-rich regions, often associated with the CSBs themselves, but there was no reliable pattern to the distribution of these regions, for example, near the beginning or end of genes.

**Table 2:** Conserved Sequence Block (CSB) characteristics in five newly described species of chewing lice in the parvorder Trichodectera. DNA lengths are in base pairs (bp).

| species | CSB1 length | CSB1 identical sites | CSB1 pairwise identity | Maximum distance between CSBs<sup>a</sup> | CSB2 length | CSB2 identical sites | CSB2 pairwise identity |
| --- | --- | --- | --- | --- | --- | --- | --- |
| *Geomydoecus geomydis* | 78 | 92% | 98% | 207 | 89 | 87% | 96% |
| *Geomydoecus centralis* | 89 | 96% | 99% | 192 | 98 | 91% | 97% |
| *Thomomydoecus minor* | 66 | 100% | 100% | 226 | 156 | 92% | 98% |
| *Stachiella octomaculatus* | 73 | 97% | 99% | 111 | 44 | 82% | 94% |
| *Neotrichodectes mephitis* | 291 | 84% | 95% | Only one CSB |  |  |  |

<sup>a</sup> Distance between CSB1 and CSB2 varies by chromosome and includes length heteroplasmy on some chromosomes. Reported distance includes gaps inserted for alignment.

In addition to the relatively large (291 bp) CSB found on all *N. mephitidis* chromosomes, there was an additional non-gene sequence that was common to the *nad3*, *nad6* and *trnI* circles. These sequences represented an additional 264 bp of common sequence at the end of the 291 bp CSB shared by all *N. mephitidis* chromosomes. On the *trnI* chromosome, the 264 bp of additional common sequence was interrupted by 54 bp of inserted sequence that was identical to a non-gene, non-CSB region of the *N. mephitidis atp8*/*atp6* chromosome. Thus, in *N. mephitidis*, there were shared non-coding sequences among some, but not all chromosomes in addition to the shared non-coding sequences common to all chromosomes of each species.

We also noted unexpected regions of sequence identity between portions of protein-coding genes on separate chromosomes. Within *G. geomydis*, for example, there was an 82-bp region of *cox1* that was identical in nucleotide sequence to a region near the end of the *G. geomydis nad2* gene (Fig 1). Other species shared similar patterns, wherein there was a region of sequence identity between *cox1* and *nad2*. In *G. centralis* and *T. minor*, the region of identity within each species was 81 bp in length (Figs 2 and 3). In *N. mephitidis* (Fig 4), there was a stretch of 78 bp that was identical between the two genes except for a single nucleotide. A pattern of nucleotide-sequence matching between the *cox1* and *nad2* genes did not occur in *S. octomaculatus* (Fig 5). The *cox1* and *nad2* regions that were highly similar or identical within a species were not identical among species, showing a pattern of sequence divergence consistent with taxonomic relationships.

The *trnL1* and *trnL2* genes were unexpectedly similar to one another in four species: *G. geomydis*, *G. centralis*, *T. minor*, and *S. octomaculatus*. Within each of these species, the *trnL1* and *trnL2* genes only differed from each other at one or two nucleotide positions among their 70-bp length, including the key position in the anticodon that makes the two tRNAs function differently. On the other hand, the *trnL1* and *trnL2* genes of *N*. *mephitidis* differed from one another at 12 nucleotide positions.

### Phylogenetic analysis

After aligning gene sequences for all 11 taxa and confirming alignment quality by visual inspection, we retained twelve genes for phylogenetic analysis: *atp6*, *cob*, *cox1*, *cox2*, *cox3*, *nad1*, *nad2*, *nad3*, *nad4*, *nad5*, *rrnL*, and *rrnS*. The trimmed alignments were concatenated to produce a dataset of 10,972 bp (S1 Data). Partitioned maximum-likelihood analysis yielded a well-resolved phylogeny with 84-100% bootstrap support for all ingroup nodes (Fig 6). Three *Bovicola* species formed a monophyletic group, as did the three species of *Geomydoecus*. The *Geomydoecus* species exhibited a sister group relationship with *Thomomydoecus minor*, and together these taxa formed a monophyletic group of lice that parasitize pocket gophers.

**Fig 6.**
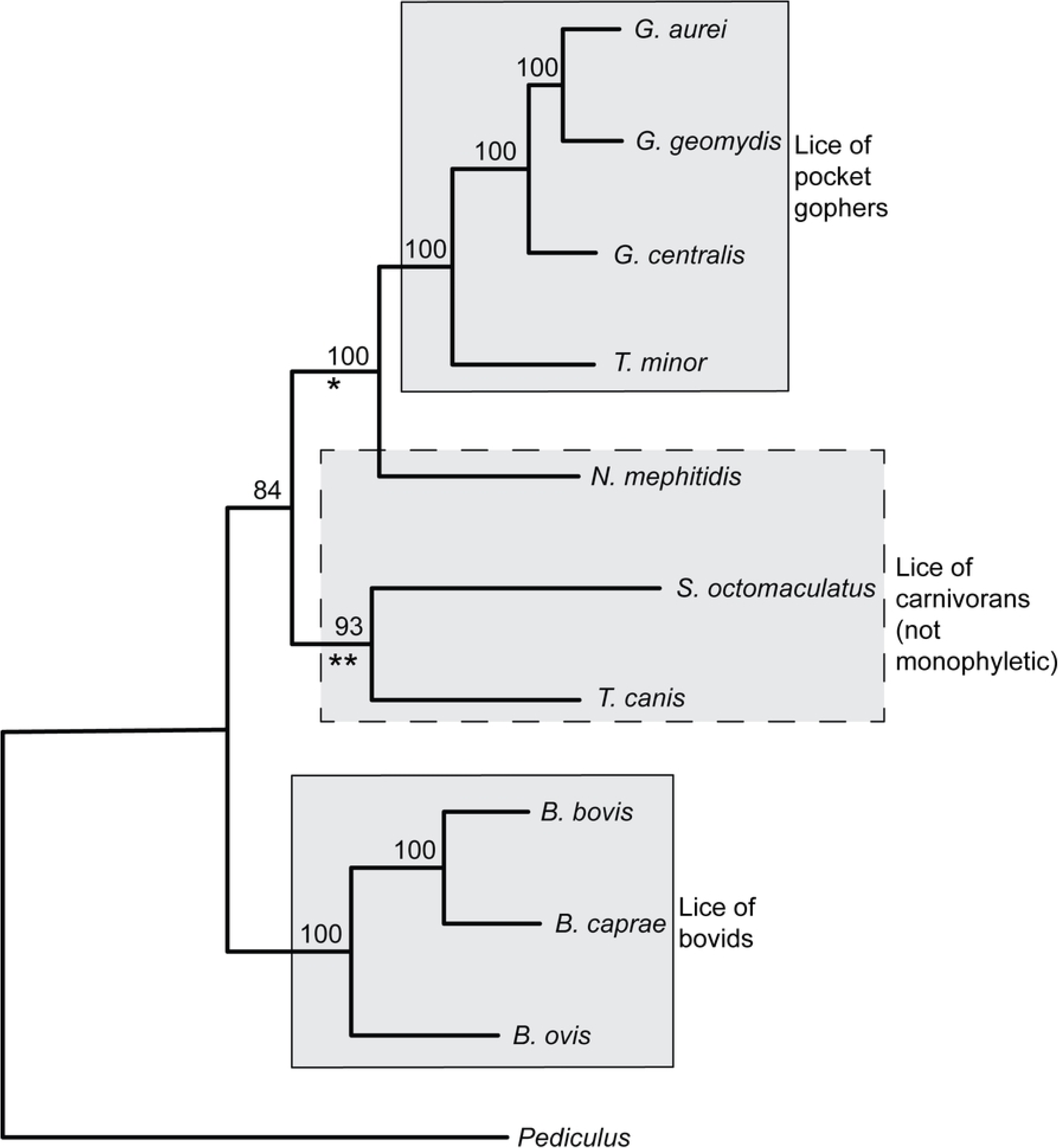
Partitioned maximum-likelihood analysis of ten species of chewing lice (Trichodectera). The human body louse, *Pediculus humanus corporis*, was used as the outgroup. Numbers represent 10,000 non-parametric bootstraps. Asterisks correspond with features discussed in text.

Interestingly, the three species of lice that parasitize carnivorans did not form a monophyletic group (Fig 6). Instead, *N. mephitidis* (the louse of skunks) was more closely related to the lice of pocket gophers than to the other lice of carnivorans; this relationship was strongly supported (100% bootstrap). Two other species of lice that parasitize carnivorans, *S. octomaculatus* and *T. canis*, appeared monophyletic (Fig 6).

## Discussion

### Chromosomal structure, size, and gene content

This work expands the number of trichodecteran lice to eleven, including the five species described for the first time herein and the six previously described [10,32–34]. All trichodecteran species studied possess mitochondrial genomes made up of 12, 13, or 14 circular minichromosomes (Table 1, [10,33,34]). This places trichodecteran genomes within the range exhibited by the closest relatives of the parvorder, Anoplura and Rhynchophthirina, which have 9-20 minichromosomes [9,11–20].

The total size of chromosomes was similar among trichodecteran species (Table 1, [10,33,34]). *Geomydoecus* and *Thomomydoecus* lice are understood to be sister genera [38], and among the four species examined in these genera, chromosome sizes tended to be similar. Differences in total size of chromosomes among close relatives were the result of differences in both non-gene content and inferred gene size (Table 1, [10,33,34]). For most chromosomes, *B. ovis* exhibited the largest chromosome in nearly every comparison among species, primarily due to the substantial amount of non-gene content on the chromosomes of this species (e.g., 1,014 bp on the *cox3* chromosome [10]). In contrast, *S. octomaculatus* typically had the smallest chromosome among species with comparable gene content, due largely to its smaller non-gene region (e.g., 267 bp on the *nad5* chromosome; Table 1).

Among the trichodecterans studied thus far, the number of genes per chromosome and the relative locations of the largest genes are the same among species with the notable exception of the *nad3* gene. This gene occurs in a variety of chromosomal locations in the trichodecterans, including alone on a chromosome in *N. mephitis* and *B. caprae*, with *cob* in *B. bovis* and *B. ovis*, and with *nad6* in the three *Geomydoecus* species, *T. minor*, *S. octomaculatus*, *T. canis*, and *T. melis* (Table 1, [10,33,34]). Although *nad3* has been found on the *nad6* chromosome in seven trichodecterans, the relative location of *nad3* differed between the four lice of pocket gophers vs. *S. octomaculatus*, *T. canis,* and *T. melis*, suggesting that the co-occurrence of these genes is not homologous in the two groups. Mapping gene location on trichodecteran phylogeny (Fig 6) suggests a multi-step history of gene arrangement with multiple fusion and/or fission events with respect to the *nad3* gene. Chromosome maps of a larger number of trichodecteran species would be needed for a clearer assessment of chromosomal fusion and fission events, but it is apparent even from this limited phylogenetic analysis that the history of gene arrangement, and rearrangement, is a complicated one. Moreover, studies of both chewing lice and sucking lice have demonstrated that the *nad3* gene can be found in a variety of locations, either alongside other protein-coding genes or in isolation [10,12–16,18–20], suggesting that *nad3*, the smallest of the mitochondrial protein-coding genes, readily shifts among genomic neighborhoods in a diverse group of species with fragmented genomes.

Even though most rRNA and protein-coding genes appear to be relatively stable in their position on separate chromosomes throughout the history of the trichodecterans, tRNA genes have been somewhat more mobile. Of the 22 tRNA genes, 13 were always associated with a particular protein-coding gene, but nine were found in alternative locations in different species (Table 1, [10,33,34]). The relative mobility of tRNA genes has also been reported in the fragmented mitochondrial genomes of the anopluran and rhynchophthirinan relatives of trichodecteran lice [9,11–20].

The reversed direction of transcription of *trnT* that was observed in *S. octomaculatus* is shared by *T. canis*, *B. bovis*, *B. ovis*, and *B. caprae* (Table 1, [10]). Mapping this character onto the inferred phylogeny of trichodecteran lice (Fig 6) suggests that the reversed orientation is ancestral for the group and that *trnT* was reoriented to the forward direction in the common ancestor of *N. mephitidis* and the pocket-gopher lice (* on Fig 6). The orientation of *trnT* is also variable among members of the parvorders Anoplura and Rhynchophthirina that have been studied [9,11–20], suggesting that the orientation of this gene is flexible in lice.

### Conserved sequence blocks

Within the non-gene region of each chromosome, all trichodecterans studied exhibited CSBs (Table 2, [10,33,34]). Members of the parvorders Anoplura and Rhynchophthirina also exhibit CSBs, as do the more distantly related nonparasitic booklice with fragmented mitochondrial genomes [7,9,11–20]. In a fragmented mitochondrial genome, each chromosome must bear a region of DNA sequence that would drive its own replication and the transcription of its genes. The CSBs examined in trichodecteran lice are predicted to form stable secondary structures and therefore may be important to the initiation of one or both of these processes. Given the fundamental importance of transcription and replication, natural selection could demand some level of constraint in the nucleotide sequence of these regions such that they can form appropriate secondary structure. One of the more dramatic examples of sequence conservation among CSBs examined here can be seen in *T. minor*, which exhibited a 66-bp region that was identical among all 12 chromosomes (Table 2). This strong sequence identity may be partially explained by the action of purifying selection.

However, purifying selection may not be the main driver of sequence similarity among minichromosomes. First, CSB sequences we examined were unique to each species studied and differed too much for credible nucleotide sequence alignment in any of the trichodecterans, even among sister taxa such as *G. aurei* and *G. geomydis*. The conservation of CSBs within individuals, coupled with relatively large differences in CSBs among species, suggests that sequences within a species are actively maintained as an outcome of gene conversion, whereby mitochondrial minichromosomes with different genes align and recombine, resulting in the reinforcement of the common sequence identity of these regions [7,9,10,12–16,18–20,57].

In many anopluran lice, AT-rich sequences have been reported near the beginning of gene clusters with GC-rich sequences near the ends of genes, presumably corresponding with start and stop of transcription [11–15,17,19,20]. However, Dunn and Shao [27] analyzed the transcription products of mitochondrial genomes of human lice and found that AT-rich sites were not transcribed at significantly higher rates than the regions upstream of them. This suggests that transcription may begin at multiple locations on the chromosome rather than only at the start of gene clusters. In contrast, GC-rich sequences at the ends of gene clusters acted more consistently as transcription stop sites, terminating a significant portion of transcripts [27]. In the trichodecteran species we studied, GC-rich and AT-rich regions were often found in the CSBs themselves, but we observed no discernable pattern in the positions of GC-rich sites, AT-rich sites, or CSB’s relative to the beginnings or ends of gene clusters, so we hypothesize that transcription may begin at multiple sites on trichodecteran chromosomes as well. Transcript data for trichodecteran lice would allow a direct test of which regions function as transcription start and stop sites in this group.

### Other evidence of recombination

In addition to the remarkable sequence identity of CSB regions described above, which may signal ongoing recombination events, our analysis identified four features of the newly described genomes that exhibited unexpected within-individual nucleotide-sequence identity among chromosomes. Our PCR-based approach, which amplified large portions of each chromosome and analyzed multiple reactions and many clones per chromosome with long regions of overlapping sequence, makes artifactual assembly an unlikely explanation for these patterns. We therefore attribute the unexpected similarity of these four regions on different chromosomes to recombination events, as has been proposed for other fragmented mitochondrial genomes [7,9,12,15,20,21].

First, in *N. mephitidis* we found evidence of potential recombination among the *nad3*, *nad6* and *trnI* chromosomes, where an identical 264-bp non-gene sequence (separate from the 291-bp CSB) occurs only on these three chromosomes. Although *nad3* and *nad6* occur together in some trichodecterans, in *N. mephitidis,* these occur on separate chromosomes. The *trnI* gene is found with *cox1* in all trichodecterans except *N. mephitidis*, where it is on a separate chromosome with no protein-coding genes (Table 1, [10,33,34]). Therefore, it is unlikely that *trnI* would have shared a chromosome with *nad3* or *nad6* ancestrally, meaning that the 264-bp sequence that these three chromosomes share is better explained by recombination among chromosomes that were already fragmented than by duplication of a *nad3/nad6* chromosome followed by degradation of the redundant genes. A second area of unexpected similarity occurred on the *N. mephitidis trnI* chromosome, where the 264-bp non-gene sequence was interrupted by a 54-bp insertion identical to non-gene, non-CSB sequence found only on the *atp8*/*atp6* chromosome. This structure is hard to explain by simple duplication and is evidence of actual exchange between chromosomes. The identical nature of these sequences, undifferentiated from one another by mutation, is consistent with ongoing recombination among chromosome-specific non-gene regions.

A third example of unexpected sequence identity occurred between protein-coding genes on different chromosomes. Five trichodecteran species, including the four pocket-gopher lice and *N. mephitidis*, each exhibited a 78-82 bp region of identical sequence in *cox1* and *nad2* (Figs 1-4; *N. mephitidis* differs at a single nucleotide). We found no such similarity between *cox1* and *nad2* in *S. octomaculatus* (Fig 5), *T. canis*, *B. ovis*, *B. bovis*, or *B. caprae* [10]. Mapping the presence of shared *cox1* and *nad2* sequence onto our reconstructed phylogeny indicates a single origin of this feature (* on Fig 6). Alignment of the shared region from all five species showed differentiation that corresponds with phylogenetic history, from about 85% identity between the sister taxa *G. aurei* and *G. geomydis* to 73% between *N. mephitidis* and its nearest relatives. Because the sequences are identical, or nearly so, within each species but differ between species, we infer that an ancient recombination event between the two genes in the common ancestor of *N. mephitidis*, *Geomydoecus*, and *Thomomydoecus* gave rise to the shared region, which then diverged in separate lineages. Within each species, the identical regions of cox1 and nad2 indicate that differences are being removed faster than they accumulate by mutation, which is most parsimoniously explained by ongoing recombination or gene conversion between chromosomes. Our data do not resolve how the homologous sequence first came to occupy both chromosomes (for example, through duplication or insertion), only that ongoing exchange between the two genes is the most likely explanation for the pattern we see.

We also observed surprising similarity of the *trnL1* and *trnL2* genes in six trichodecteran species, including *G. aurei*, *G. geomydis*, *G. centralis*, *T. minor*, *T. canis*, and *S. octomaculatus*, where these genes reside on different chromosomes (Table 1, [10,33,34]). In each of the six species, the two genes differed from each other at only one or two nucleotide positions among their 70-bp length, including the key position in the anticodon that makes the two tRNAs function differently. This pattern of similarity was not shared by *B. caprae*, *B. ovis*, and *B. bovis*, in which the two genes differed from each other by 27-41 nucleotides, consistent with independent evolution of the two genes. The *trnL1* and *trnL2* genes of *N*. *mephitidis* differ from one another by 12 nucleotides, an intermediate amount of gene similarity. A few anopluran lice also share nearly identical *trnL1* and *trnL2* [9,12,17], while others do not [11,13,19,20]. It is possible that *trnL1* and *trnL2* genes were homogenized once by gene duplication or recombination in the ancestor of a monophyletic group of all carnivoran- and pocket-gopher lice (Fig 6), with subsequent recombination continuing to homogenize these two genes, albeit less completely in *N. mephitidis* than in the other six species. Selection presumably maintains the anticodon difference, since recombination alone would erase it. Alternatively, separate instances of recombination-driven similarity may have arisen independently, which seems likely given that both anoplurans and trichodecterans include species with nearly identical genes and species without.

Taken together, these examples of similar or identical sequences occurring on different chromosomes are consistent with the conclusion that recombination has acted on both gene and non-gene regions of trichodecteran mitochondrial genomes. Previous researchers have noted presumed recombination-driven similarities between protein-coding genes and/or tRNA genes within the genomes of anopluran sucking lice. In those species, the region of similarity varies widely in length, from as few as 20 to as many as 127 nucleotides [9,12,15,20]; the 78-82 bp *cox1*/*nad2* regions we report fall within that range. Patterns of nucleotide-sequence sharing among chromosomes have been demonstrated in the parvorders Anoplura [9,12,15,20], Rhynchophthirina [18], and Trichodectera (herein), as well as more distantly related lineages with fragmented mitochondrial genomes [7,21], indicating that recombination among chromosomes may be a general feature of fragmented mitochondrial genomes in lice, not a peculiarity of certain groups.

### Phylogenetic Analysis

Twelve mitochondrial protein-coding and rRNA genes yielded robust alignments. Only the highly variable *atp8*, *nad4L*, and *nad6* genes failed to yield usable alignments. These three genes were singled out by Lin et al. [58], who demonstrated that they are under positive selection (rather than purifying selection) in a diverse representation of four parvorders of phthirapteran lice, and we note that these appear to be the most rapidly evolving genes in trichodecteran mitochondrial genomes as assessed by low (64-70%) sequence identity even between sister taxa, *G. aurei* and *G. geomydis*.

Phylogenetic analysis indicated some relationships that would be expected based on taxonomy and previous work. The three *Bovicola* species, which parasitize ungulates, formed a monophyletic group, with *B. caprae* and *B. bovis* appearing as sister taxa as they did in previous analyses [10]. Three species of *Geomydoecus* appeared as a monophyletic group that was closely related to *Thomomydoecus minor*, yielding a monophyletic group of lice that parasitize pocket gophers, a conclusion that is consistent with previous work based on morphology [38]. Pocket gophers and their chewing lice have served as a model system for cospeciation, with many host and parasite groups showing significant phylogenetic congruence [43,59–63]. However, three of the four species included in this study were collected from the same species of host, *T. bottae.* These species therefore represent a departure from strict host-parasite cospeciation.

Interestingly, the three species of lice that parasitize carnivorans did not form a monophyletic group (Fig 6). Instead, *N. mephitidis* (the louse of striped skunks) exhibited a more recent evolutionary relationship with the lice of pocket gophers rather than with the other two species of lice from carnivorans (Fig 6). The sister relationship of *N. mephitidis* to the chewing lice of pocket gophers that was demonstrated by nucleotide sequences (* on Fig 6) was also supported by two unique gene arrangements: 1) the presence of *trnP* on the *cox3* chromosome, and 2) the transcription of *trnT* on the same strand as other coding- and tRNA-genes instead of in “reverse” orientation on the opposite strand as it is transcribed in *S. octomaculatus* (Table 1) and other trichodecterans [10,33]. Species in this clade also each had an identical sequence region located in both *cox1* and *nad2* (Figs 1-4).

Other carnivoran lice included in the analysis, *T. canis* [10] and *S. octomaculatus*, were monophyletic with 93% bootstrap support (** on Fig 6). This relationship was supported by four unique gene arrangements that suggested a shared evolutionary history for these two species, including 1) the lack of *trnY* on the *cox2* chromosome, 2) the presence of *trnL1* on the *cox3* chromosome, 3) the unique arrangement of tRNA genes and the presence of *nad3* on the *nad6* chromosome, and 4) the presence of a tRNA-only chromosome with *trnY* and *trnP* on it (Table 1).

The paraphyletic nature of carnivoran lice reported here was previously noted by Johnson et al. [64] who used genomic data to show that *N. mephitidis* is more closely related to a pocket gopher louse (*G. aurei*) than either is to another carnivoran louse, *Stachiella larseni*, found on mink [64]. These authors used cophylogenetic reconstruction to show that pocket gophers acquired their trichodecteran lice from carnivorans [64]. This host-switch was also posited in a cospeciation analysis by Taylor and Purvis [65]. Because pocket gophers and their trichodecteran lice have a history of widespread cospeciation [43,59–63], the host-switch from carnivore to rodent must have predated much of the diversification among the geomyid rodents and among their trichodecteran lice on them. Johnson et al. [66] discussed this host switch in the context of observations that raptors may have transferred lice to a prey species through unsuccessful predatory interactions. We add that this transfer of lice from predator to prey could also have occurred through temporary, shared occupancy of the burrow of a pocket gopher species that was ancestral to extant pocket-gopher genera. A passive transfer of lice in the past is plausible given that modern skunks and long-tailed weasels, both hosts of *Neotrichodectes* species, are known to enter the burrows of pocket gophers [67–69], thus facilitating host switching through indirect contact. Indeed, pocket gopher lice (*Geomydoecus*) have been discovered in low numbers as “stragglers” on the long-tailed weasel, *Mustela frenata* [70]. It would be interesting to know if all species of *Neotrichodectes* are more closely related to pocket gopher lice than they are to other carnivoran lice, or if the genus is paraphyletic. A more robust sampling of *Neotrichodectes* and their hosts could provide a better understanding of the degree of cospeciation within this group and perhaps provide interesting perspective on the history of host switching and cospeciation.

## Acknowledgements

For assistance in the laboratory, we are grateful to many undergraduate students at the University of Northern Iowa, including Aubree Blanco, Skye Franklin, Payton Harle, Jaylie Hicklin, Jessica Lopez-Clemente, Joshua Mixdorf, Jordan Rubner, Aaliyah Streif, Claire Wade, and Anna Wood. The authors acknowledge the support of the Freiburg Galaxy Team: Person X and Björn Grüning, Bioinformatics, University of Freiburg (Germany), funded by the German Federal Ministry of Education and Research BMFTR grant 031 A538A de.NBI-RBC and the Ministry of Science, Research and the Arts Baden-Württemberg (MWK) within the framework of LIBIS/de.NBI Freiburg.

## Supporting Information

S1 Table: Primers used to generate mitochondrial sequences of *Geomydoecus geomydis*.

S2 Table: Primers used to sequence the mitochondrial genome of *Geomydoecus centralis*.

S3 Table: Primers used to generate mitochondrial sequences of *Thomomydoecus minor*.

S4 Table: Primers used to generate mitochondrial sequences of *Neotrichodectes mephitidis*.

S5 Table: Primers used to generate mitochondrial sequences of *Stachiella octomaculatus*.

S1 Data: Nexus file showing twelve genes that were aligned, trimmed, and concatenated for phylogenetic analysis.

